# Hobotnica: exploring molecular signature quality

**DOI:** 10.1101/2021.09.12.459931

**Authors:** Alexey Stupnikov, Alexey Sizykh, Alexander Favorov, Bahman Afsari, Sarah Wheelan, Luigi Marchionni, Yulia A. Medvedeva

## Abstract

A Molecular Features Set (MFS), is a result of vast diversity of bioinformatics pipelines. In case when MFS is used for further analysis to distinguish between phenotypes, it is often referred to as a signature. Lack of the “gold standard” for most experimental data modalities makes it hard to provide valid estimation for a particular MFS’s quality. Yet, this goal can partially be achieved by analyzing inner-sample Distance Matrix (DM) and their power to distinguish between phenotypes.

The quality of a DM can be assessed by summarizing its power to quantify the differences of inner-phenotype and outer-phenotype distances. This estimation of the DM quality can be construed as a measure of the MFS’s quality.

Here we propose **Hobotnica**, an approach to estimate MFS’s quality by their ability to stratify data, and assign them significance scores, that allows for collating various signatures and comparing their quality for contrasting groups.

## Introduction

A signature based on a predefined Molecular Features Set (MFS), which is designed to distinguish biological conditions or phenotypes from each other — is one of major concepts of bioinformatics and precision medicine. In this context, signatures typically originate from MFS from contrasting experimental data of two or more sample groups, which differ phenotypically. These MFS incorporate information on the differences between the groups. The nature of the MFS depends on the modality of the original data. For instance, the MFS provided by the Differential Gene Expression approach is a list of Differentially Expressed Genes (DEG); Differential Methylation analysis provides Differentially Methylated Cytosines or Regions (DMC and DMR) as MFS, etc.

A significant number of mutational, expression and methylation-based signatures have recently been published and they are actively used in Research and Transnational Medicine. Examples of expression-based signatures involve genesets for clinical prognosis (e.g. PAM50 (***Parker et al. (2009)***), MammaPrint (***Cardoso et al. (2016)***) for Breast Cancer), for pathways and gene enrichment analysis (e.g. MsigDB collections (***Subramanian et al. (2005)***)), for drug re-purposing (e.g. LINCS project(***Liu et al. (2015)***)).

Direct quality assessment for MFS is currently hardly possible, since there are no’gold standard’ datasets where active Molecular Features are explicitly known. In this manuscript, we propose a novel approach - **Hobotnica** - that allows for measurement of MFS quality by addressing the key property of the signature, namely, its quality for data stratification.

Hobotnica leverages the quality of Distance Matrices obtained from any source in order to assess quality of the MFS from any data modality compared to a random MFS. In this study, we demonstrate its application on transcriptomic signatures.

## Results

### Approach

The Hobotnica approach is as follows: For a given data set *W* and a given Molecular Features Set (*S*) we derive the inter-sample distance matrix (*DM*(*S,W*)). Then we assess the quality of *DM* (and, thus, of *S*) with a summarizing function (*α*(*DM*(*S*)) = *α*(*DM*(*S*),*Y*) or by abuse of notation *α*(*DM*(*S*))) where (*Y*) represents the labels of samples.

We desire the function *α* to gauge if the inner-class samples are closer to each other than to outer-class samples. If no difference exists from one class to another, *α* must be close to zero and as the difference grows, *α* grows. In ideal case of a perfect separation, *α* reaches its maximum at 1:

- *α* ∈ [0, 1]
- *α* → 1 ⇔ High groups stratification quality
- *α* → 0 ⇔ Low groups stratification quality

Under the Null hypothesis of Hobotnica ((*H*_0_)), no significant difference exists between *α*(*S*) and the *α* of an equal-sized general random set. On the contrary, the Alternative (*H_A_*) hypothesizes that *S* generates higher *α* than most random *S*′ of the same size. To estimate a Null distribution for *Hobotnica′s α*, we applied a permutation test. As our default options, we use Kendall distance as the distance measure and Mann-Whitney-Wilcoxon test as the summarizing function.

### Validation

To validate our approach in the first case study we extracted RNA-seq expression dataset for Prostate Cancer from TCGA on counts level(***Rahman et al. (2015)***). As Molecular Feature Sets we recruited C2 collection of molecular signatures from MSigDB (***Subramanian et al. (2005), Liberzon et al. (2011)***) that contains a number of Prostate-related genesets. For the second case study we took PAM50 molecular signature, designed for various Breast Cancer types classification, and applied it to several datasets (***Marusyk et al. (2016)***)(***Daemen et al. (2013)***)(***Costello et al. (2014)***)(***Rahman et al. (2015)***)(***Varley et al. (2014)***). In both cases, the counts were normalised to cpm. For each geneset H-score and its p-value with BH correction were computed.

Prostate-related C2 genesets clearly demonstrated highest values of H-score and sufficient statistical significance(Fig.1.A), as well as data stratification (Fig.1.B), which is expected for Prostate Cancer vs Control contrast. Genesets not attributed to Prostate Cancer related processes did not achieve statistical significant p-values. (Table1).

**Figure 1.**
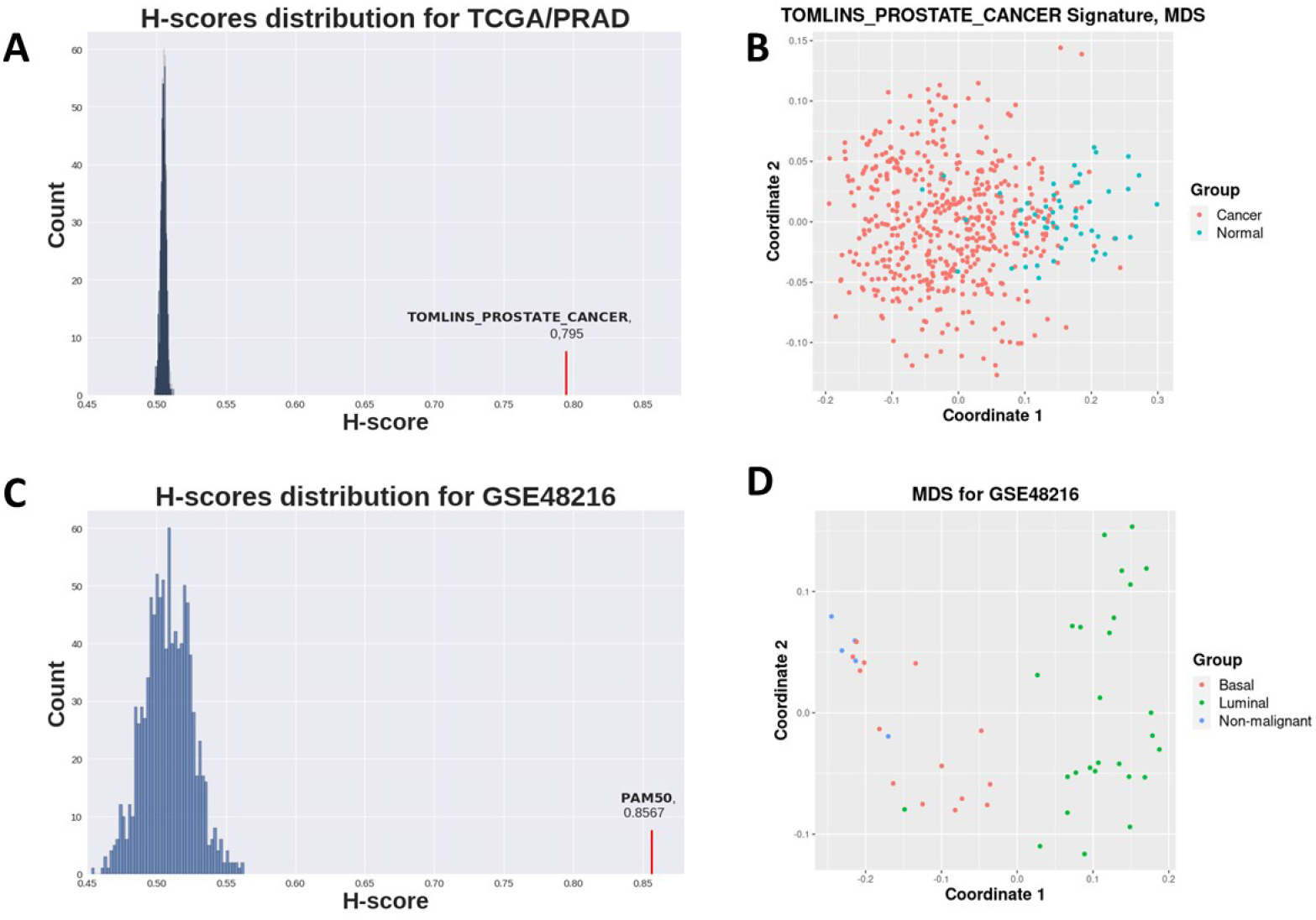
A: Distribution of H-scores for random genesets (blue) on TCGA Prostate Cancer vs Normal dataset (see Tab.1) and Tomlins prostate geneset H-score (red). B: MDS for TCGA Prostate demonstrates samples separation with Tomlins geneset. C: Distribution of H-scores for random genesets (blue) on GSE48216 Breast Cancer dataset (see Tab.2) and PAM50 geneset H-score (red). D: MDS for GSE48216 Breast Cancer dataset samples separation with PAM50 geneset.

**Table 1.**
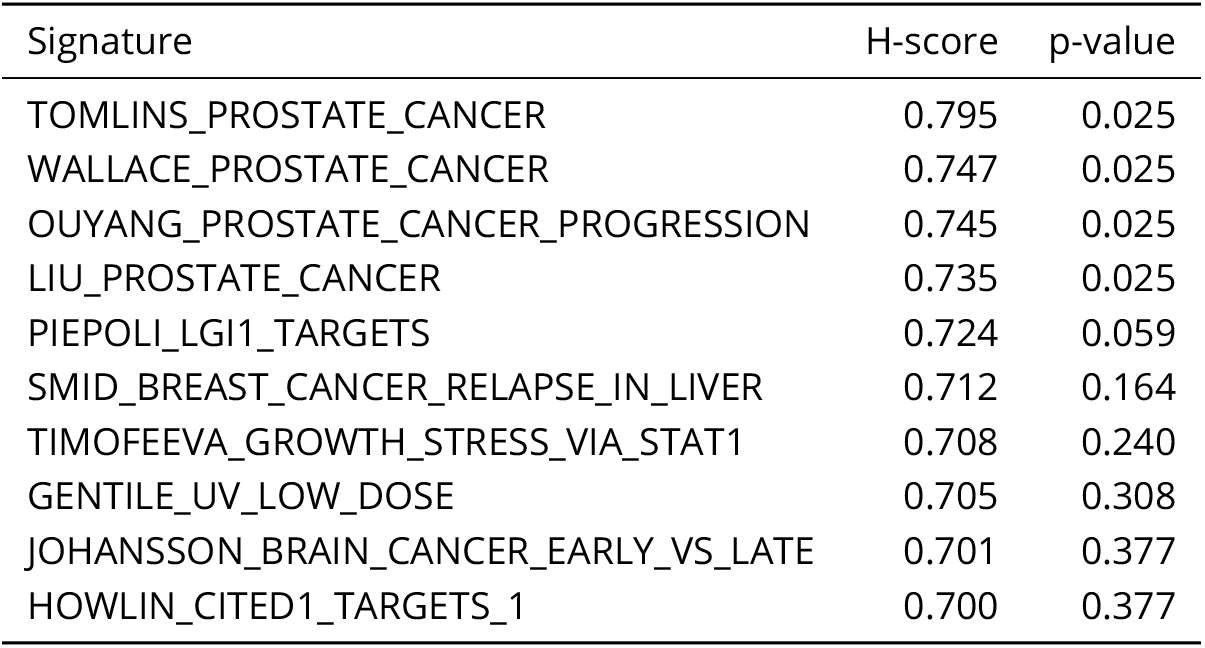
10 C2-CGP Gene Signatures with highest H-scores

PAM50 signature evidently separates samples in for GSE48216 dataset (Fig.1.C). H-scores for random genesets for the same dataset are significantly lower than an H-score for PAM50 (Fig.1.D). Clearly, PAM50 signature demonstrates high quality of stratification for the samples of various Breast Cancer datasets with high H-score values and statistically significant p-values (Table 2).

**Table 2.** PAM50 results

| GEO Accession | Sample size | Groups in dataset | H-score | p-value |
| --- | --- | --- | --- | --- |
| GSE58135 | 168 | 6 | 0.772 | 7e-4 |
| GSE62944 | 1067 | 5 | 0.8892 | 0.0003 |
| GSE48216 | 46 | 3 | 0.8567 | 0.0003 |
| GSE80333 | 10 | 3 | 0.9765 | 0.0003 |

Thus, in the first case study, Prostate Cancer related genesets from C2 collection, when applied to Prostate Cancer dataset, delivered highest H-scores and most significant and p-values proved to demonstrate best scores and performance. Likewise, in the second case study, PAM50 expression signature applied to several heterogeneous Breast Cancer datasets delivered high H-score values along with significant scores of p-values.

### Application

An important question that researches often face is establishing the optimal size of the retrieved signature. The exact number of genes to be retrieved from the set of all significant genes is an important parameter that is essential for signature’s application. To explore the optimal size of DE signature we performed Hobotnica analysis for top DE p-value ordered gene signatures of various lengths. For the reference we performed DGE analysis for Breast Cancer vs Control TCGA dataset (***Rahman et al. (2015)***) with DESeq2 (***Love et al. (2014)***) and edgeR (***McCarthy et al. (2012)***). Top 100 genes for each method were retrieved, as well as genes with highest variance in expression.

H-scores for every signature then were computed (Fig.2.A.). For this dataset DESeq2 provided a signature with the highest quality score. Then, we calculated H-score for signatures of various lengths (Fig.2.B.) Surprisingly, the signature quality is non-monotonously dependant on the signature length, i.e. number of genes in the signature. The pattern also varies for DE models. Additionally, for the best-performing model, DESeq2, the a signature quality is generally declining with the length. Thus, increasing number of genes in a signature may not improve its quality, and an optimal gene signature length for the DE analysis result may be established: DESeq2 signature reaches its maximum H-score at 13 genes and edgeR at 11 in this case.

**Figure 2.**
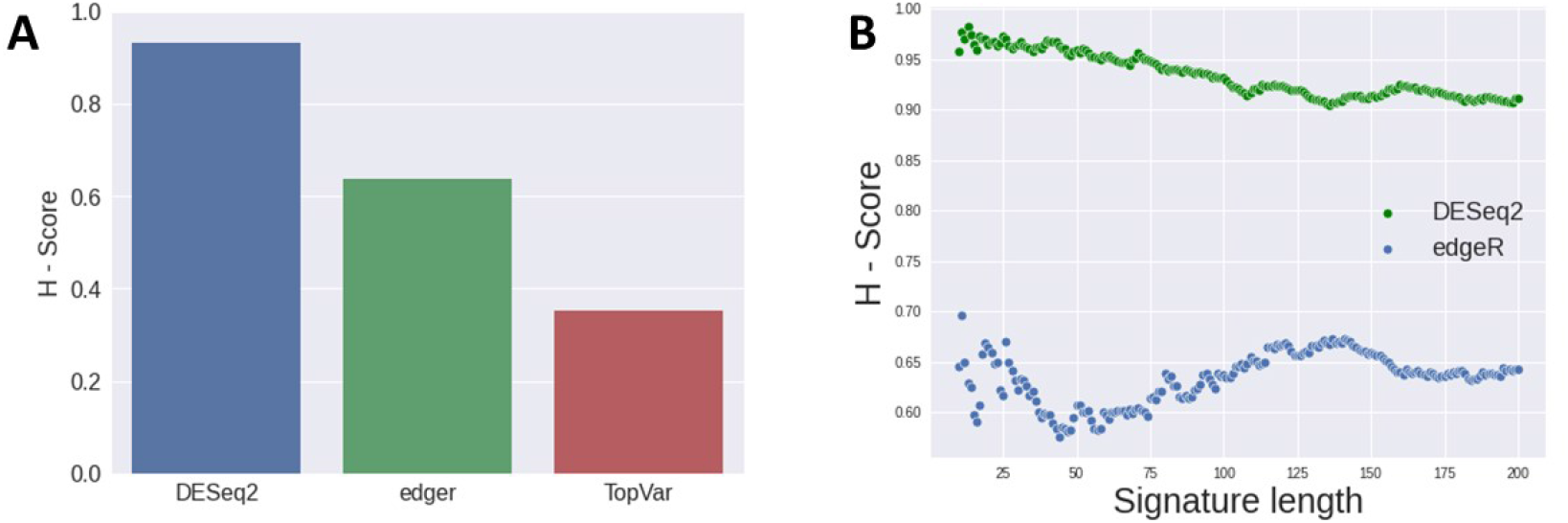
A: H-scores delivered by top 100 gene signatures from various DE models applied to TCGA Breast Cancer data. B: Change of H-score with the length of gene signature derived from DESeq2 and edgeR models

## Discussion

*Hobotnica* is designed to quantitatively evaluate Molecular Feature Set’s quality by their ability for data stratification from their inter-sample distance matrices, and to assess the statistical significance of the results. We demonstrated that Hobotnica can efficiently estimate the quality of a Molecular Signature in the context of Expression data.

Suggested method can be used to evaluate Molecular Feature sets of various nature: retrieved in DGE, Differential Methylation analysis, Mutation/SNV calling or Pathways analysis, as well as data modalities from other types of Differential Problem. In addition, assessing H-score values for various lengths of the same set or signature will help with its structure optimization, which may be especially important in clinical applications.

*Hobotnica* is available as an R package at https://github.com/lab-medvedeva/Hobotnica-main

## Methods and Materials

### Problem formalization

If a Molecular Feature Set (*S*), that presumably incorporates information on the contrast between groups of samples with known samples annotation *Y* in Data *D* is in place (*H : S*), we can compute Distance Matrix between samples *DM* (*f*(*S*|*D*) → *DM*) and then introduce a measure *α* (*g*(*DM*|*Y*) → *α*) of signature quality for Data *D* stratification.

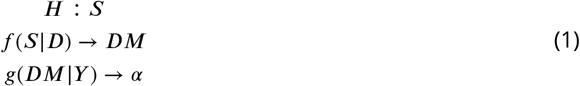

When instead of a single *GS* a set of hypotheses {*H*_1_ : *GS*_1_,*H*_2_ : *GS*_2_,…,*H_n_* : *GS_n_*} is in place, for each Gene Signature *GS_i_* corresponding Distance Matrix *DM_i_*, can be generated, and than, in turn, particular value of the measure *α_i_*:

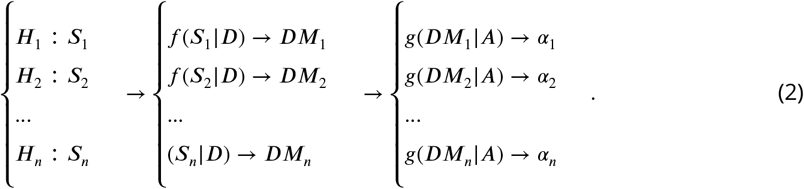

Thus, for every MFS *S,* from set of hypotheses {*H*_1_ : *S*_1_,*H*_2_ : *S*_2_,…,*H_n_* : *S_n_*} H-score *α_i_*, maybe computed, resulting in a set 〈*α*_1_,*α*_2_,…*α_n_*〉. Comparing *α* values allows for corresponding Feature Sets qualities ranking and selecting the most informative Signatures for the Data *D*.

To assess statistical significance of each obtained H-score *α_i_*, we compute empirical *p-value* via generating a distribution of H-scores for set of random MFS.

### Availability

We implemented Hobotnica as an R package available at https://github.com/lab-medvedeva/Hobotnica-main https://github.com/lab-medvedeva/Hobotnica-main. It contains an implementation of the Hobotnica measure, statistical analysis for significance, and several auxiliary functions for visualizing results and parallel processing.

## Acknowledgements

We thank Frank Emmert-Streib, Leslie Cope and Elana Fertig for fruitful discussions. The study was supported by Ministry of Science and Higher Education of the Russian Federation (agreement no. 075-15-2020-899) and by the NIH grants R01DE027809 and P30CA006973.

