## Supplemental File 1 for "Hobotnica: exploring molecular signature quality"

### Hobotnica: A Technical Note

Alexey Stupnikov

May 2021

#### Introduction

Hobotnica is an algorithm that can be used to assess the quality of a Molecular Feature Set, or Molecular Signature. It uses the experimental design (i.e. information for each sample to which group it belongs) and a signature-induced samples distance matrix. The introduced measure summarises between-samples relationship structure. The tool is implemented and available as an R package at https://github.com/lab-medvedeva/Hobotnica-main

#### Motivation

A Molecular Features Set (MFS), is a result of vast diversity of bioinformatics pipelines. In case when MFS is used for further analysis to distinguish between phenotype, the analysis is often referred to as a signature. Usually, molecular signatures are considered a solution for a Differential Problem, that implies contrasting two or more cohorts of samples with different phenotypic properties.

Due to various reasons differential analysis of experimental data with different bioinformatical models results in sets of molecular signatures, which often differ in structure and content. Lack of "gold standard" data for most experimental data modalities makes it impossible to provide any hard evidence in favor of any particular molecular signature.

We propose and mathematical model and an approach based on rank statistics to introduce a measure, that will allow to estimate molecular signature's quality and, therefore to compare signatures in a set of candidate Molecular Feature Sets.

#### Formalization

Consider a candidate Molecular Feature Set ($S$) and its corresponding inter-point distance matrix ($D(S)$) for samples in the data set. Hobotnica requires a summarizing function ($\alpha(S) = \alpha(D(S), Y)$) where ($Y$) represents the label of samples. Summarizing function must gauge if the inner-class samples are closer relative to outer-class samples. In ideal case, if there is no difference, the summarizing function is close to zero and as the difference grows the function increases. Under the Null ($H_0$), Hobotnica Hypothesizes that $S$ makes no difference in $\alpha(S)$ and the Alternative ($H_A$) Hypothesizes that $S$ increases $\alpha(S)$ compared to other sets with the same size.

#### Model

When we have computed inter-point distance matrix for a Molecular Feature Set $D(S)$, we require $\alpha(D(S), Y)$ (or by abuse of notation $\alpha(S)$). The intuition regarding this metric is as follows:

- $\alpha \in [0, 1]$
- $\alpha \to 1 \Leftrightarrow$ High outer-class distance relative to inner-class distance
- $\alpha \to 0 \Leftrightarrow$ Low outer-class distance relative to inner-class distance

As our default options, we use Kendall distance as the distance measure and Mann-Whitney-Wilcoxon test as the summarizing function.

For two groups of samples, $R$ and $G$, taking into account the nature of Distance Matrix (0-diagonal and symmetry), corresponding lower triangle matrix may be considered. The distance values of this matrix $d_{pq}$ can be ranked, producing new matrix with natural values $r_{pq}$ (Fig.1).

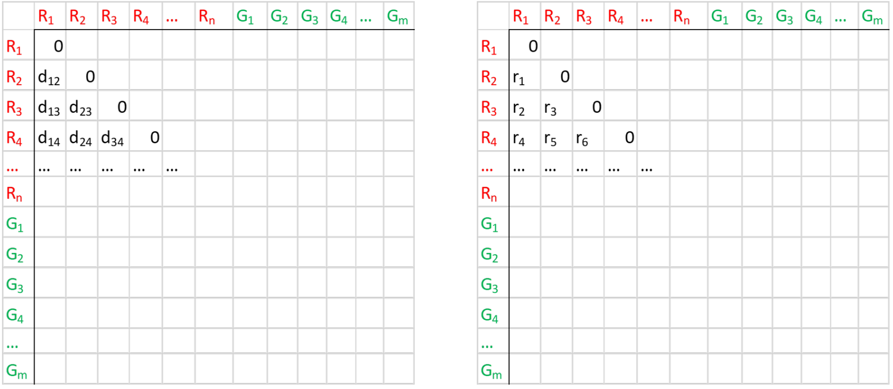


Figure 1: Distances ranking procedure: each distcance element $d_{pq}$ is substituted with corresponding rank value $r_{pq}$

Subseting the matrix to values corresponding to in-class distances (Fig.2), we can compute sum of these ranks $\Sigma$. This sum's values correspond to various samples relationship situations: the lower the sum is, the closer samples from the same class to each other, the more accurately Distance Matrix structure corresponds to the samples labels.

Given the numbers of samples is $n$ in group 1 and in $m$ group 2, total number of ranks (and therefore max value of a rank) in un-subsetted triangle matrix will be

$$M = \frac{(m+n)(m+n-1)}{2} \tag{1}$$

Likewise, total number of ranks is subsetted to in-class values will be

$$N = \frac{(m^2+n^2-(m+n))}{2} \tag{2}$$

$\Sigma$ reaches its min value $A$ if the subset procedure selects minimal values of ranks, or, in other words, ranks in the selected part of rank matrix are values from 1 to $M$:

$$A = \sum_{i=1}^{M} i \tag{3}$$

The minimal value is reached when the lowest ranks reside in the diagonal squares of the distance matrix that correspond to in-group distances, and, therefore, best groups separation. Similarly, max value $B$ is delivered when maximal values of Ranks Matrix are selected - from $(N-M)$ to $N$:

$$B = \sum_{i=(N-M)}^{N} i \tag{4}$$

Thus, $\Sigma \in [A, B]$. Now performing a scaling of compact $[A, B]$ to $[0, 1]$ finally allows us to retrieve value $\alpha$ with requested properties:

$$F : [A, B] \to [0, 1], F(A) = 1, F(B) = 0 \tag{5}$$

$$F : S \to \alpha \tag{6}$$

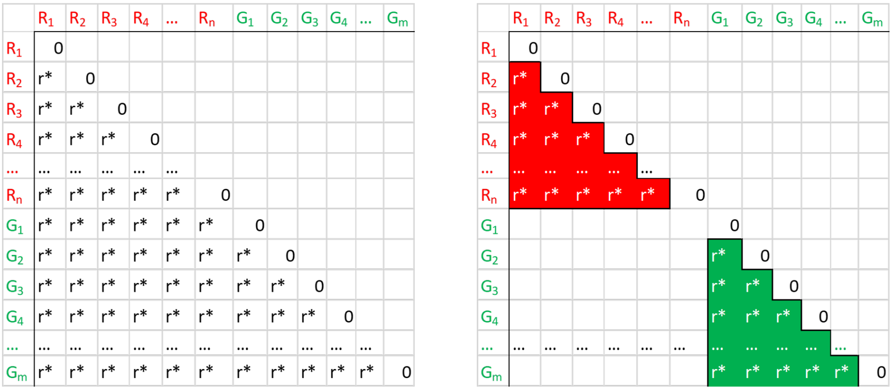


Figure 2: Subsetting in-class ranks

This procedure finally allows to introduce measure $\alpha$ with necessary listed properties. We refer to this measure as $\boldsymbol{Hobotnica}, or \underline{H-score}$.

Thus, for every Gene Signature $GS_i$ from set of hypotheses $\{H_1 : GS_1, H_2 : GS_2, ..., H_n : GS_n\}$ H-score $\alpha_i$ may be computed, resulting in a set $\langle \alpha_1, \alpha_2, ...\alpha_n, \rangle$. Comparing and ranking $\alpha$ values allows for corresponding Gene Signatures qualities ranking and comparison.

We use ranking matrix instead of the distance one for several reasons. First of all, we want to make our algorithm less dependent on distance choice. With ranking, theoretical minimal and maximal values of sum for subsetted in-class matrix can be calculated analytically. In addition, ranking helps to combat the negative effects of outliers.

### Statistical inference

To assess statistical significance of the obtained H-score $\alpha$ we compute empirical p-value via generating a distribution of H-scores for set of random Gene Signatures.

The p-value is computed as follows. A large number of signatures (with the same lengths as signatures in the hypotheses set) is sampled with replacement from the list of all Molecular Features in the dataset (e.g. transcripts), thus resulting in a collection of random signatures, i.e. lists consisting of randomly selected Molecular Features (transcripts in case of expression data). This yields an empirical distribution of H-scores for the dataset. An example of such distribution is displayed in Fig.3

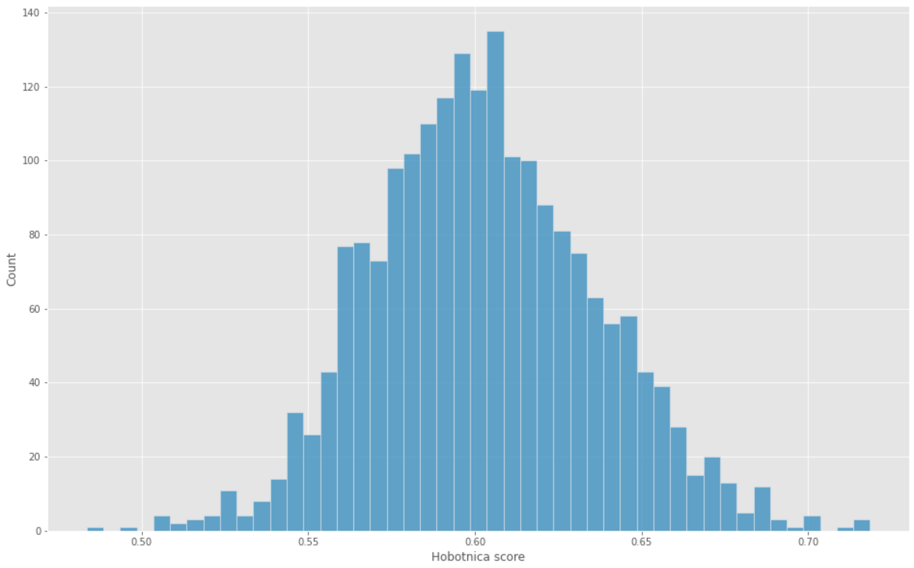


Figure 3: An example of H-scores distribution for random signatures

With the distribution in place, the p-value of a given signature H-score is calculated as the fraction of random signatures with greater or equal H-score. For numeric stability, we add a pseudo-count to the numerator.

### Pseudocode

**Algorithm 1** Hobotnica algorithm

**procedure** HOBOTNICA($distMatrix, annotation$)
▷ distMatrix is a matrix of distances between samples
▷ annotation describes grouping of samples
$rankMatrix \leftarrow ranked\, distMatrix$
$innerSum \leftarrow 0$
$uniqueGroups \leftarrow a\, vector\, with\, unique\, groups\, from\, annotation$
$lengths \leftarrow empty\, array\, of\, length\, equal\, Length(uniqueGroups)$
**for** $i \leftarrow 1, Length(uniqueGroups)$ **do**
$group \leftarrow uniqueGroups[i]$
$lengths[i] \leftarrow number\, of\, samples\, in\, group\, i$
$temporaryInclassSum \leftarrow$ in-group ranks sum
$innerSum \leftarrow innerSum + temporaryInclasSum$
**end for**
$l \leftarrow Length(annotation)$
$biggesRank \leftarrow \frac{l(l-1)}{2}$
$nInclassElements \leftarrow \frac{\sum_j lengths[j](lengths[j]-1)}{2}$
$minSum \leftarrow nInclassElements(nInclassElements + 1)$
$maxSum \leftarrow nInclassElements(2 \cdot biggesRank - nInclassElements + 1)$
$normalizationFactor \leftarrow maxSum - minSum$
**return** $1 - \frac{innerSum - minSum}{normalizationFactor}$
**end procedure**
